# Motor “laziness” constrains fixation selection in real-world tasks

**DOI:** 10.1101/2023.02.13.528190

**Authors:** Charlie S. Burlingham, Naveen Sendhilnathan, Oleg Komogortsev, T. Scott Murdison, Michael J. Proulx

## Abstract

People coordinate their eye, head, and body movements to gather information from a dynamic environment while maximizing reward and minimizing biomechanical and energetic costs. Such natural behavior is not possible in a laboratory setting where the head and body are usually restrained and the tasks and stimuli used often lack ecological validity. Therefore, it’s unclear to what extent principles of fixation selection derived from lab studies, such as inhibition-of-return (IOR), apply in a real-world setting. To address this gap, participants performed nine real-world tasks, including driving, grocery shopping, and building a lego set, while wearing a mobile eye tracker (169 recordings; 26.6 hours). Surprisingly, spatial and temporal IOR were absent in all tasks. Instead, participants most often returned to what they just viewed, and saccade latencies were shorter preceding return than forward saccades. We hypothesized that participants minimize the time their eyes spend in an eccentric position to conserve eye and head motor effort. Correspondingly, we observed center biases in the distributions of fixation location and duration, relative to the head’s orientation. A model that generates scanpaths by randomly sampling these distributions reproduced the spatial and temporal return phenomena seen in the data, including distinct 3-fixation sequences for forward versus return saccades. The amount of the orbit used in each task traded off with fixation duration, as if both incur costs in the same space. Conservation of effort (“laziness”) explains all these behaviors, demonstrating that motor costs shape how people extract and act on relevant visual information from the environment.

**Significance Statement:** Humans display remarkably precise yet flexible control of eye and body movements, allowing for a wide range of activities. However, most studies of gaze behavior use the same setup: a head-restrained participant views small images on a computer. Such lab studies find that people avoid looking at the same thing twice, and hesitate in cases when they do. We had people perform nine everyday activities while wearing glasses with embedded eye tracking, and surprisingly found that they did the opposite, often returning to what they just viewed and expediting these “return” eye movements over others. A tendency to keep the eyes centered in the head, which we speculate helps to conserve motor effort, explained these behaviors for all tasks.

## 1 Introduction

Fixation selection can be described as an active inference process which maximizes reward while minimizing costs for the agent. In this framework, gaze shifts and maintenance of fixation both serve to reduce uncertainty about task-relevant properties of the environment, preventing sub-optimal actions [1–4]. As an example, fixating an ambiguous road sign while driving enables reading and heeding the sign’s text, “Detour”, reducing the total time taken to reach a destination (a type of reward). And because active inference takes place in a physical body, each muscular contraction and neural computation involved in generating a gaze shift, fixation, or body movement incurs specific costs. Therefore, the brain must estimate these costs and account for them when planning actions, balancing them with expected rewards and reductions in uncertainty. Indeed, simply assuming that the brain optimizes metabolic energy when moving can, on its own, explain a number of un-intuitive motor behaviors during locomotion [5]. And task- and motor-dependent cost functions in optimal feedback control models have proven successful in predicting behavior for other motor systems; for example, those involved in reaching [6]. Likewise, the idea that the brain optimizes energy required for neural computations, an important constraint in frameworks like efficient coding and resource rationality, can explain a large range of cognitive and perceptual phenomena [7–10].

Decades of research on fixation selection and gaze behavior in psychology and neuroscience has generated many important insights, including the discovery of reward sensitivity of neural oculomotor circuitry, as well as the conceptual advances of saliency maps, inhibition-of-return, reinforcement learning, and active inference in understanding and modelling gaze behavior [2–4, 11, 12]. Perhaps the most general properties of fixation behavior are (1) the center bias, the tendency for the eye to remain mostly near the head orientation / center of the orbit, and return phenomena like (2) spatial inhibition-of-return, the tendency for observers to more often gaze at new locations than recently visited ones, and (3) temporal inhibition-of-return, longer fixation durations preceding a return than gazing at a new location. Some have speculated about whether biomechanical and energetic efficiency in operating the oculomotor muscles may give rise to the center bias [13–15], but there is no work to our knowledge which actually quantifies how big of an influence this has, versus other factors. Likewise, return phenomena are generally understood to arise from high-level perceptual and cognitive processes [16]; for example, the idea that inhibiting return reduces redundancy in information gathered over time [11, 17].

In this study, we measured gaze in nine real-world tasks while observers wore a mobile eye tracker and quantified return phenomena and center biases in each. We found that participants returned more often than expected in all tasks and displayed shorter fixations preceding return than forward saccades; that is, opposite to past findings taken as support for spatial and temporal inhibition-of-return. We observed large eccentricity-dependent differences in fixation probability and duration in all tasks, which on their own quantitatively explained the probability of return as well as fixation durations preceding returns in a simple random sampling model. Our findings suggest that an eccentricity-dependent variable strongly contributes to return phenomena. We speculate that this variable is effort. And we propose the hypothesis that energy optimality, specifically minimization of motor costs associated with expected eye and head movements, can parsimoniously account for all of our observations. Consistent with this hypothesis, the amount of the orbit used in a given task traded off almost linearly with average fixation duration, as if both incur costs in the same space.

## 2 Results

Each participant performed a subset of nine real-world tasks while their gaze was recorded in head-centered coordinates using the Tobii Pro Glasses 2 mobile eye tracker (accuracy, ±0.62^*°*^, temporal sampling rate, 50 Hz; see Methods for task and equipment details). Fixations were detected for each recording, yielding estimated scanpaths for each task and participant. We analyzed scanpaths using standard procedures from the literature designed to identify spatial and temporal inhibition-of-return. Next, we simulated synthetic scanpaths by drawing random samples from the probability distribution of fixation locations and the map of average fixation duration — both functions of eye position relative to the (approximate) head orientation. To test whether this random model could capture the phenomena seen in the data, we applied the same analyses to the data and model scanpaths.

### 2.1 Fixation position and duration displayed task-dependent center biases

Past studies have identified a center bias in fixation position, where “center” corresponds with the center of the field-of-view (FOV) in head-fixed free-viewing of images, or the camera/head orientation in head-unrestrained experiments [14, 15, 18, 19]. Likewise, it is known that more eccentric fixations are maintained for a shorter duration, and that more central fixations are maintained for a longer duration [13]. We characterized both of these phenomena for each task in our data by estimating (1) the probability distribution of fixation position, and (2) a map of average fixation duration, both in head-centered coordinates. To summarize the statistics of fixation position across observers, we divided the FOV into 60 × 60 equal size bins, and summed the total number of fixations across observers and tasks (Fig. 1). Approximately 95% of fixations were within the central 35^*°*^of both the horizontal and vertical conditional density passing through the origin. Approximately 50% were within the central 13^*°*^, vertically, and the central 8^*°*^, horizontally. Thus, the pooled distribution was considerably peakier than a Gaussian (i.e., leptokurtic), and narrower horizontally than vertically. Note that many eccentric bins were empty for all tasks and participants, that is, participants very rarely or never fixated in the far periphery. We observed strong center biases in each task, but with large inter-task differences in the center and dispersion of the position distribution (Fig S1). For fixation duration, we estimated the mean fixation duration in milliseconds within each spatial bin, and for visualization purposes plotted only bins containing more than 10 fixations, for otherwise the mean was dominated by sampling error. For the pooled data (Fig 1), we observed a similar shape to the fixation position distribution, but more dispersed. The fixations closest to the head orientation were around 800 ms, whereas the most eccentric fixations were around 150 ms. There was large inter-task variability in the offset of the duration map (i.e., some tasks had longer fixations at each position), as well as in its center and shape (Fig. S2). But generally, fixations were shorter the more eccentric they were.

**Fig. 1.**
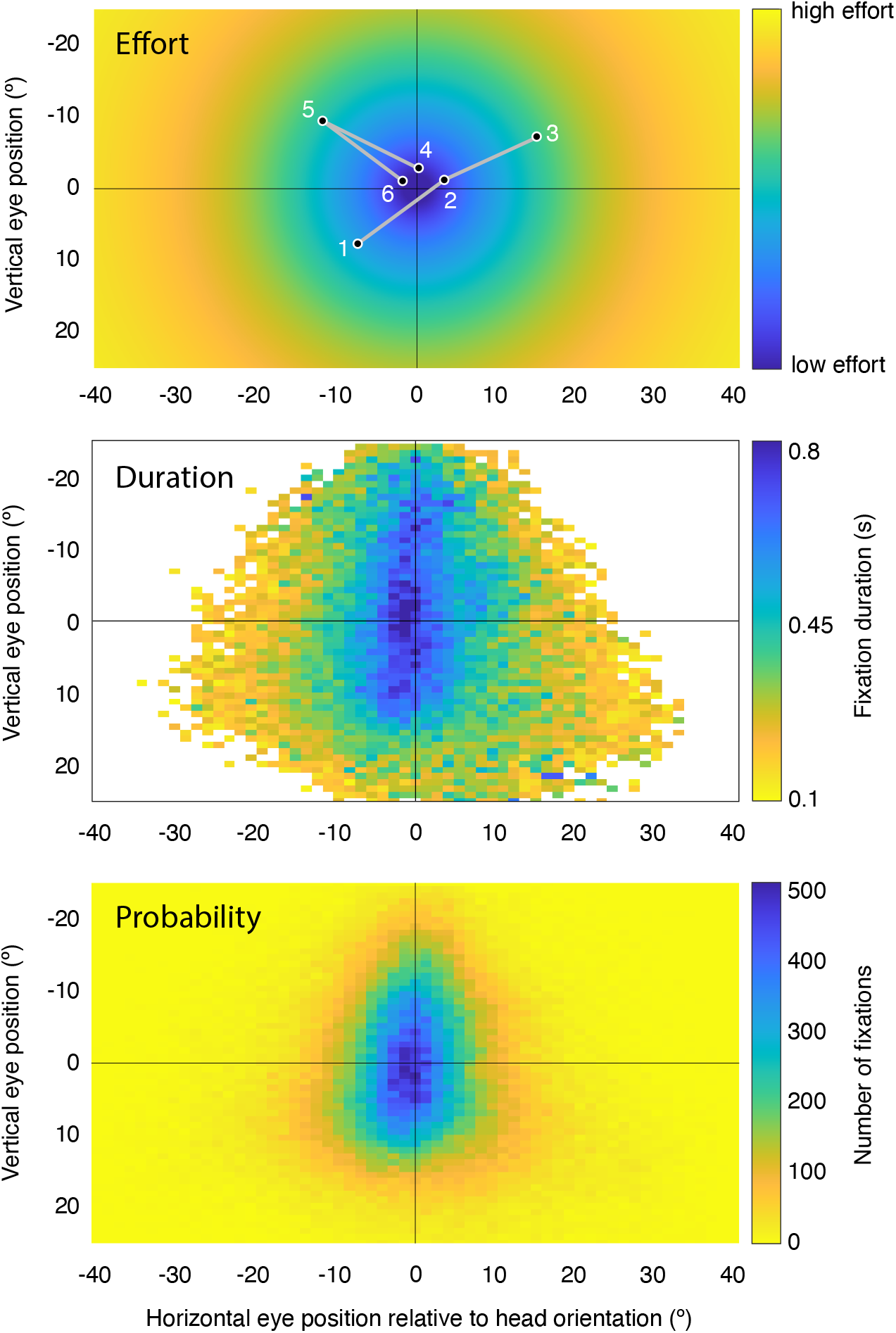
Effort minimization hypothesis of fixation selection. Top panel, cartoon depicting effort required to fixate a given orbital angle (cross-hairs represent head orientation, which is close to the center of the orbit). We hypothesize that observers orient their head toward an object of interest, such that when the eye is in the primary position (0,0), it is roughly oriented at the object. Fixations 1, 2, and 3 in the cartoon scanpath represent a forward saccade, and 4, 5, and 6 represent a re-fixation. The hypothesis predicts that people will make fewer saccades to locations far from the center of the orbit (validated with data in middle panel), and that when they do fixate far from the center of the orbit, they will linger there for a shorter amount of time (validated in bottom panel). The middle panel depicts the average fixation duration across all tasks and recordings and the bottom panel depicts the total number of fixations in a given location in the field-of-view. Note that the hypothesis predicts that distinct temporal sequences will arise for forward and return saccades - fixation 2 in the sequence is predicted to be longer than fixation 5. Given that more eccentric fixations are rarer and forward sequences take up more space than return sequences, the second fixation in a forward sequence is more likely to be closer to the center of the orbit than the second fixation in a return sequence. To generate a synthetic scanpath, we draw a sample from the positional distribution in the bottom panel to simulate a single fixation, then look up its duration in that bin in the middle panel, and then repeat. White values in the middle panel represent positions for which there were fewer than 10 fixations, such that the mean was dominated by sampling error.

### 2.2 Real-world scanpaths do not exhibit spatial inhibition-of-return — observers re-fixate often in all tasks

To characterize spatial inhibition-of-return in our data, we estimated the joint distribution of relative saccade (log) amplitudes and directions (i.e., the angular difference between successive saccades), and measured the ratio of the number of forward vs re-fixation saccade sequences by looking at the frequency of saccades within a 90^*°*^ slice centered on either 0^*°*^ or 180^*°*^, respectively, from the previous (following [17]; see Methods for our operational definition of a saccade). Surprisingly, we observed a much larger number of re-fixations relative to forward saccades in all tasks (Fig. 2A, Fig. S3, Fig. S8B), with the ratios of re-fixations to forward saccades ranging from 1.5-3.5 across tasks. An extreme example of this was the browsing task (ratio = 3.5), when observers scrolled on their phones and read text, making frequent saccades up to the previous visual field location (where new text appeared) and then moving their eyes down as they read.

**Fig. 2.**
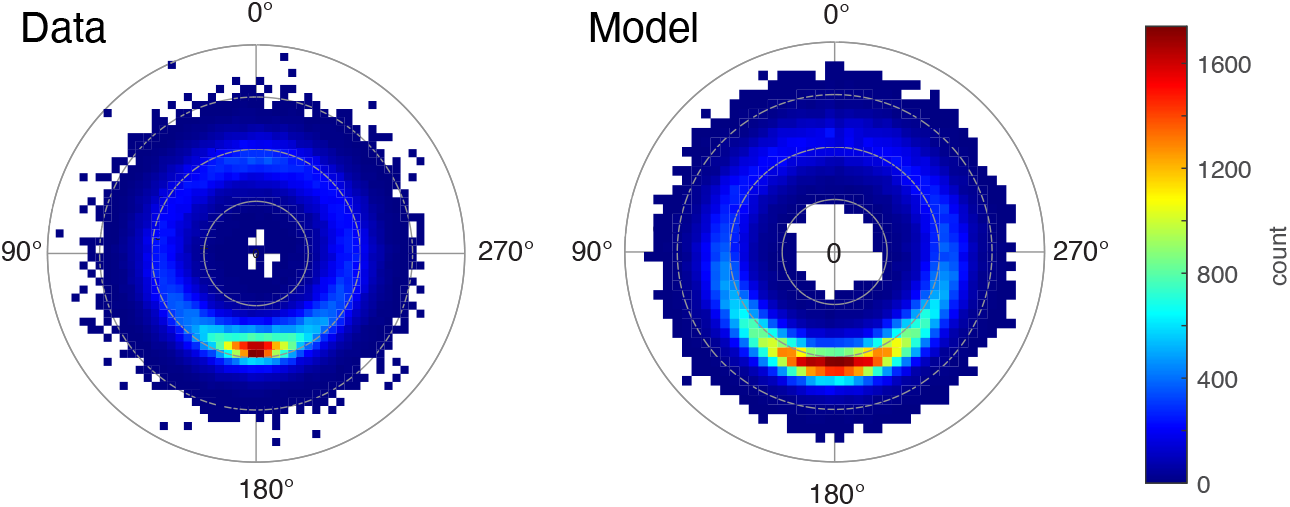
Human and model scanpaths exhibit high rates of re-fixation. Left panel, relative saccade amplitude and direction joint distribution, for data pooled over all tasks and participants. Second ring out from the center corresponds to an amplitude difference of 0^*°*^. That is, the most probable type of sequence was a precise return saccade. Radius axis is in log space, and its limits were scaled per subplot. Right, same format but for simulated scanpaths from the model (N is matched in data and model). The ratio of re-fixations to forward saccades in the model was 4.38 (vs. 2.44 in the data). This demonstrates that sampling randomly from the distribution of fixation locations is sufficient to generate re-fixation rates comparable to, but slightly overestimating, that seen in the data.

### 2.3 A random fixation selection model reproduces high re-fixation rates

Given that the distribution of fixation positions was focused near the head orientation, it follows that even for a participant who selects fixations randomly and with temporal independence from this distribution, re-fixations should be more probable than forward saccades (assuming that the total amplitudes are roughly matched). This is simply because forward sequences take up more of the orbit than re-fixations of the same total amplitude — that is, the average endpoints are more eccentric — and because eccentric fixations are rarer (see Fig. 1A. for illustration). To test this idea, we simulated scanpaths for each task separately and overall across all tasks by drawing random samples from the corresponding distribution of fixation positions. The amount of synthetic and real data was matched. We then estimated the joint distributions of relative saccade directions and amplitudes for the synthetic data, for each task, and computed the spatial IOR ratios as well. The model reproduced the predominance of re-fixations seen in the data in all tasks (Fig. 2A, Fig. S3). Overall, the spatial IOR ratio was around two times higher in the model than in the data, and the model did not capture the task dependency in this ratio seen in the data (Fig. S8B). This is because, in the model, forward saccades are equally improbable in each task whereas, in the data, their probability varies more across tasks (and is slightly higher overall), which drives task differences in the spatial IOR ratio.

### 2.4 Fixation durations were longer preceding forward than return saccades, the opposite of temporal IOR

Temporal inhibition-of-return is the tendency for observers to fixate longer preceding a return than forward saccade, when successive saccades are matched in amplitude. Following [17], we defined forward and return saccades as 3-fixation sequences with an absolute relative amplitude of less than the 25% quantile of the distribution (|Δ*r*| *<* 25%), and a relative angle within 50^*°*^ (360^*°*^ / 7 bins) of either precisely forward (0^*°*^) or backward (±180^*°*^) [17]. The relative amplitude thresholding was done separately for each recording, given that the relative amplitudes varied between tasks (Fig. 3, Fig S3). The grand mean fixation duration across all tasks and participants was 525 ms for the second fixation in a forward sequence and 466 ms for the second fixation in a return sequence, that is, the opposite of temporal IOR (p *<* 0.001, one-tailed permutation test, 1000 permutations). We found the same pattern (or no evidence for a statistical difference) in all nine tasks (Fig. S4; one-tailed permutation tests, forward *>* return: p = 0.001, 0.011, 0.001, 0.478, 0.038, 0.28, 0.0529, 0.004, 0.265; order of tasks: Browsing, Desk, Driving, Grocery, Lego, Navigation, Restaurant, Pokémon, Walk). No task showed evidence for temporal IOR (i.e., p > 0.95, equivalent to the return *>* forward one-tailed test).

**Fig. 3.**
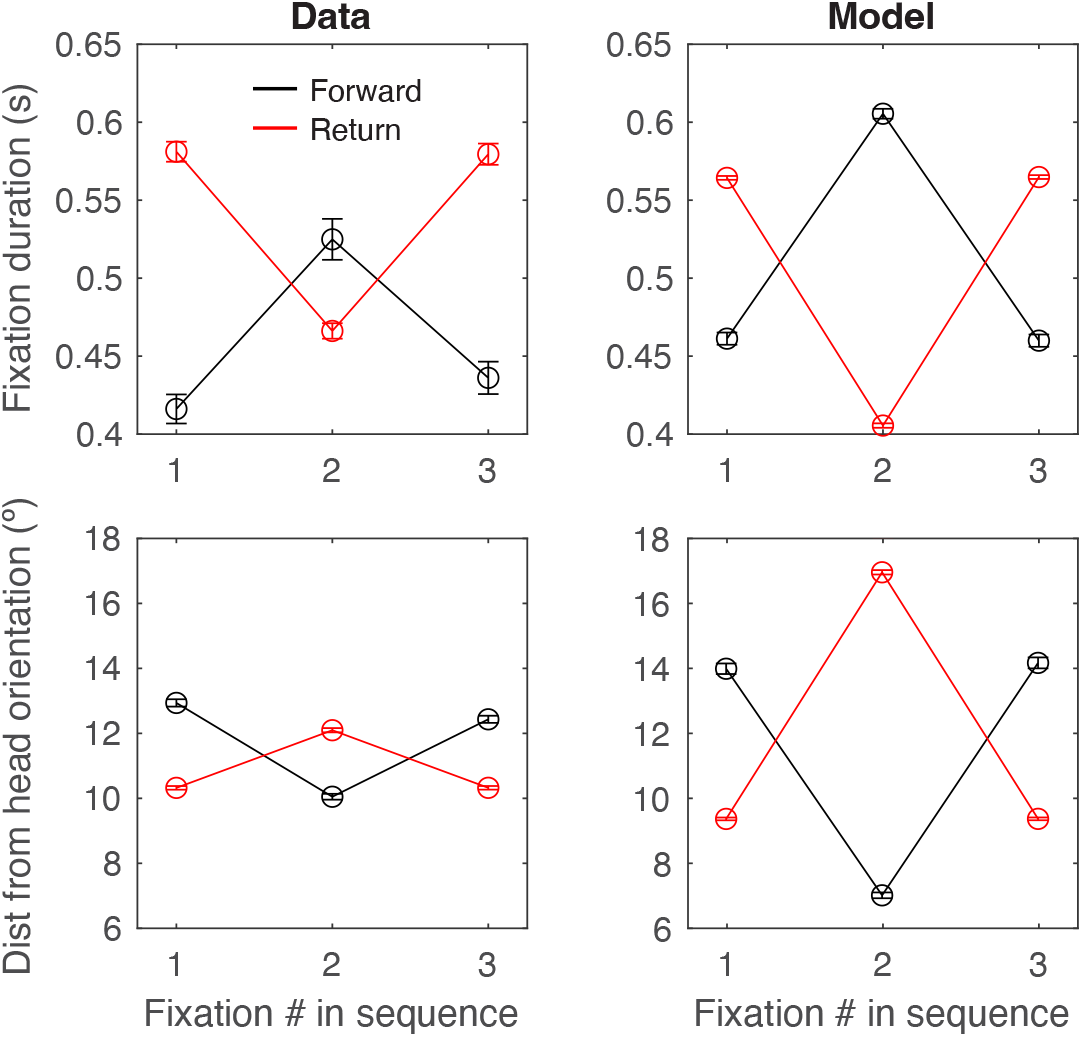
Distinct temporal sequences for forward and return saccade sequences are explained by position of the eye in the orbit, and our model predicts these sequences. Top left panel, avg. fixation duration in seconds for forward and return saccade sequences. Errorbar, SEM across recordings. A forward sequence is defined as any two saccades whose angles are within +-25^*°*^ of one another and whose amplitudes are similar (within 25% of the distribution of relative amplitude distribution for that recording). A return sequence is defined as any two saccades whose angles are within +-25^*°*^ of 180^*°*^ from one another and whose amplitudes are similar (within 25% of the distribution of relative amplitude distribution for that recording). Bottom left panel, same as top panel but the ordinate displays avg. euclidean distance from the head orientation (center of the orbit) in degrees of visual angle. Right panels, same plots but for model. In both data and model panels, N fixations = 16,008 for return sequences and N = 1,838 for forward sequences, for each datapoint plotted).

### 2.5 Forward and return saccades exhibit distinct sequences of fixation duration, which are inversely proportional to distance from the head orientation

To better understand our finding of longer fixations preceding a forward versus a return saccade, we looked at the duration of the first and third fixations in each three-fixation sequence as well. For forward sequences, observers tended to make a shorter fixation followed by a longer fixation, followed by another shorter one. And for return sequences, observers did the opposite. These distinct patterns were observed in the pooled data (Fig. 3) and in each task separately (Fig. S4). Interestingly, when we instead analyzed the distance of these fixations from the head orientation / center of orbit, we found the same patterns but nearly exactly inverted, for the pooled data and for each task separately (Fig. 3, Fig. S5). The strikingly strong inverse relationship observed here and in Fig. 1 is consistent with a mediating relationship, that is, in which the eccentricity of a fixation determines how long it is maintained for.

### 2.6 Random fixation selection reproduces fixation duration and distance sequences for forward and return saccades

To test whether the time-integrated statistics of fixation duration and position could capture the temporal dependencies we observed in scanpaths for forward and return saccade sequences, we applied the same analyses used on the data to the synthetic scanpaths we generated previously. The model exhibited longer fixations preceding forward than return saccades, as well as distinct sequences of fixation duration and distance from the center of the orbit, which matched the data in their patterns (Fig. 3). The model overestimated the difference in duration between forward and return saccades for the second fixation in the sequence, but slightly underestimated the difference for the first and third fixation — more consistent with each curve undergoing an additive offset with opposite sign, rather than a scaling. Interestingly, this means that the difference between the model and data — that is, what the model doesn’t capture quantitatively in the data — is a temporal IOR effect similar to what past studies have observed, with longer fixations preceding a return than forward saccade. This opens the possibility that our data could be described by the sum of two effects, one solely based distance from the center of the orbit (and captured by our model’s assumptions), and a second, distance-independent effect that is consistent with what has been described previously as temporal IOR, with the distance-dependent effect being considerably larger in our data than the distance-independent / temporal IOR effect.

### 2.7 Temporal inhibition-of-return is largely absent even when controlling for a fixation’s distance from the head orientation

To test the idea that a temporal-IOR-like effect may be present in our data, but obscured by a larger distance-dependent effect, we performed a distance-matched controlled analysis. If this idea is correct, we should observe longer fixations preceding a return than forward saccade, for those fixations which lie along a ring around the origin (within some tolerance) — that is, which have the same distance from the head orientation. Distance matching consisted of finding sequences that were within 2^*°*^ of the median of the distribution of distances (i.e., where there was the most data). Distance matching was done for each participant and task separately, after which we pooled the data for this analysis. We found that, on the contrary, fixation duration was statistically indistinguishable for return and forward sequences, when distance-matched (Fig. S6, permutation test, p = 0.67, N permutations = 1000). Given that the sample size after distance-matching was far less, to ensure a fair comparison, we resampled from the raw data using the same sample sizes from the distance-matched data. Specifically we resampled the data 500 times and performed one permutation test each. The median p-value was 0.0604 and 46% of p-values were *<* 0.05 (N permutations for each test = 1000). This demonstrates that there wasn’t strong evidence for the presence of temporal IOR in the data overall, even for distance-matched fixations. Next, we looked at each task separately (Fig. S7). Only one task, Lego, showed a statistically significant temporal IOR-like effect (two-tailed permutation test; p = 0.001). The browsing task had a significant effect like we saw in the non-distance-matched data with duration longer for forward than return sequences (two-tailed permutation test; p = 0.002). There was no evidence of a significant difference for the remaining 7 tasks, although some tasks, like Walk, showed a trend toward an IOR-like effect.

### 2.8 Fixation duration and the amount of the orbit used in a task traded off with each other

According to our effort minimization hypothesis, the total amount of time the eye is in an eccentric position reflects an underlying oculomotor cost which should be accounted for while also maximizing task-specific rewards. Because humans are fixating observers, this total time should be decomposed into two components — fixation position and fixation duration — both incurring the same type of cost. Some tasks naturally require more of the orbit (i.e., more eccentric fixation positions) to gather the same level of reward as others. In such tasks, the eye movement generation circuitry should correspondingly reduce fixation duration, which would equalize effort expended across many possible tasks. This idea predicts that orbital range and fixation duration should trade off with each other systematically across tasks. To test this, we measured the portion of the orbit used in a given task as the dispersion of the fixation position distribution (sum of the standard deviations of the horizontal and vertical marginal distributions; Fig S1). We measured the fixation duration as the grand mean fixation duration across all fixations in a given task. We observed a tight, nearly linear inverse relationship between the two quantities across tasks (Fig. 4; Pearson’s correlation: r = -0.95, p *<* 0.001). The Browsing task had the longest fixation duration (838 ms) and the smallest portion of the orbit was used (5.98^*°*^). The Walk task had the shortest fixation duration (264 ms) and the largest portion of the orbit was used (11.05^*°*^). This is consistent with the idea that the total amount of time the eye spends in an eccentric position reflects a motor cost that the brain accounts for systematically across tasks with different demands.

**Fig. 4.**
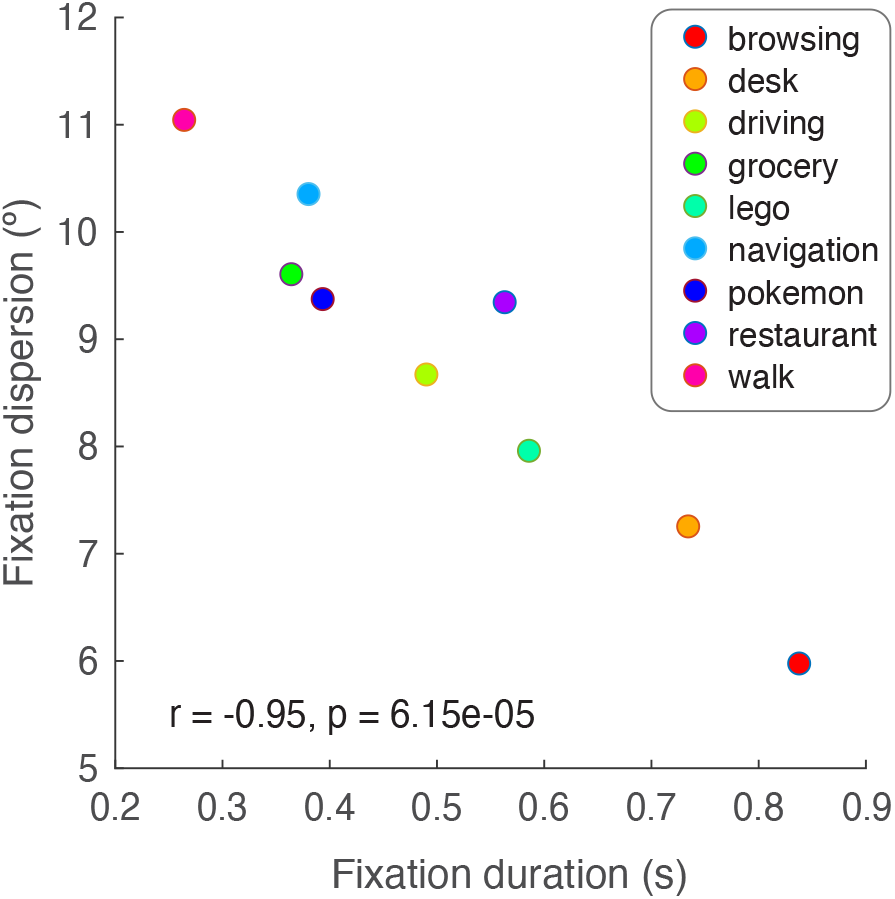
Fixation duration and the amount of the orbit used traded off with each across tasks. Left panel, grand mean fixation duration (s) for each task versus fixation dispersion (^*°*^ ; measured as the sum of the horizontal and vertical standard deviation of the distribution of fixation locations). Every color is a different task. R and p statistics are from a Pearson’s correlation.

## 3 Discussion

### 3.1 Overview

In a wide range of real-world tasks, we found that people tended to keep their eyes in a centered position and that eccentric fixations were much shorter than central ones. These tendencies explain a number of seemingly-unrelated phenomena: (1) people return to what they just viewed much more often than they view new locations, (2) fixations preceding returns are shorter than fixations preceding forward gaze shifts, and (3) there are distinct, inverted 3-fixation sequences of durations for return versus forward gaze shifts. We also found that fixation duration and orbital range traded off with each other across tasks. Our results are consistent with the idea that the perceptual-motor system conserves effort, minimizing energetic costs incurred by the specific demands of a task.

### 3.2 Spatial IOR

People re-fixate often whether the head is restrained or not. Past work has shown that the presence and extent of spatial IOR varies substantially with task and environment [2, 17, 20–22]. Broadly, tasks that incentivize exploration, like visual search and so-called “free-viewing,” have a higher ratio of forward to return saccades (i.e., they exhibit spatial IOR), whereas many other kinds of tasks exhibit the opposite (spatial *facilitation* of return) [2, 16, 17, 20]. In some tasks, like reading text on a smartphone or cooking, it’s clear *a priori* that people must re-fixate often to be able to do the task well. That said, most previous studies have used a traditional, head-restrained, static picture viewing laboratory set-up, and so it’s been unclear whether spatial IOR is present in real-world, head-unrestrained conditions [2]. A recent study, Zhang et al. 2021, compared the probability of re-fixation in a variety of tasks and viewing conditions, including head-restrained conditions while observers performed free-viewing and visual search tasks, and head-unrestrained conditions (i.e., egocentric video + mobile eye tracking) while observers either made breakfast or searched in a room for items [16]. These head-unrestrained data offer a relevant comparison for our present findings. The probability of re-fixation was around 12% on average across their datasets, being highest (30%) for the egocentric task in which observers cooked breakfast and 8% for the egocentric search task. When analyzed in the same way, the average probability of re-fixation across tasks in our data was 5% (5.4% when only analyzing periods when the head was stable, that is, *<* 3^*°*^s velocity), with the Browsing task having the largest re-fixation probability (13%). Note also, that in our data, re-fixations were the most common type of three-fixation sequence (out of all possible relative angles and amplitudes), and always more common than forward saccades (Zhang and colleagues didn’t perform this standard comparison, unfortunately). This suggests that in head-unrestrained conditions, people re-fixate exceedingly often, that is, spatial IOR is weak or absent.

### 3.3 Temporal IOR

Temporal IOR is a robust phenomenon, observed in a large number of head-restrained studies. In our study, we observed longer fixations preceding a forward saccade than a re-fixation in nearly all tasks, that is, opposite of the pattern seen in past highly-controlled lab studies [17, 20, 21]. That said, this doesn’t mean temporal IOR was absent in our observers, if considered in the original sense of a delay in information processing (along with other reductions in performance) before attending or orienting to a previously viewed target, versus other targets [11]. Rather, we admit that we can’t determine whether or not such an effect is present in our naturalistic data, but instead defer to the large body of experimental work on the topic [11]. What we can conclude is that there is a seemingly independent, eccentricity-dependent influence on fixation duration in all tasks in our dataset, which is large, and may exist alongside (e.g., sum with) temporal IOR. One piece of evidence in support of this summation account is that our model over-predicted the difference in duration between fixations preceding forward versus return saccades by about a factor of two, indicating that a second un-modelled effect with the opposite sign must be added in to reproduce the data. This second effect could be temporal IOR. The second, and weaker, piece of evidence is that when we restricted our analyses to fixations lying in a ring in head-centered coordinates (i.e., eccentricity-matched), in 2/9 tasks (Lego, Walk) we observed longer fixation durations preceding return than forward saccades — that is, consistent with past studies claiming temporal IOR.

### 3.4 Center bias

Center bias is driven, in part, by the tendency to re-center the eyes in the orbit. Observers exhibit a strong tendency to fixate the center of a computer display or picture [14, 15], or the center of the head-centered coordinate system in 360^*°*^ virtual environments [23, 24] and real environments [18, 19, 25, 26]. Past work has shown that there are a number of factors that contribute to the dispersion and center of this center bias, including task demands, environmental layout, FOV size, motor biases, photographer bias, and the tendency to re-center the eyes in the orbit [14, 15]. Likewise, in our data, we observed marked differences between tasks in the dispersion and center of the fixation position distribution. Determining the relative contribution of the multiple factors that give rise to these differences is an interesting problem, but beyond the scope of this paper. Instead, we begin from the well known observation that in head-unrestrained conditions, observers tend to roughly re-center the eyes in the orbit [27] via specific coordinated movements of the eye and head, which are well characterized [27–31]. This re-centering behavior must produce some degree of center bias. In the following sections, we will explore the factors that may contribute to this tendency to re-center the eyes in the orbit, including the mobility of the oculomotor muscles, energetic efficiency, and needs of eye-head coordination.

### 3.5 The orbital position-latency relation

Fixation duration scales inversely with the eccentricity of the eye in the orbit, with shorter fixations at more eccentric positions [13, 29, 31–34]. Fuller was the first to characterize this relation in humans while also controlling for saccade amplitude, and did so under head-restrained conditions [29]. Our head-unrestrained findings are consistent with Fuller’s, with more eccentric fixations maintained for a shorter duration in all tasks. Fuller’s explanation for the orbital position-latency relation is based on the demands of eye-head coordination during gaze shifts. When the eye is near edge of the orbit and there is target requiring a contraversive gaze shift, the ocular saccade’s onset precedes the head movement onset and ends before the head movement, leading to counter-rolling of the eye at the end of the gaze shift [13, 27, 29]. At the other extreme, when the eye is near the (other) edge of the orbit, then a target in the same direction instead requires an ipsiversive gaze shift, but the eye cannot move past the edge of the orbit so an initial head movement (with vestibular ocular reflex [VOR]) is required, followed by a delayed ocular saccade. Saccade latencies (and hence fixation durations) should be therefore longer for the second case (and for any case in-between the two extremes, like fixations near the center of the orbit), just because VOR takes some time to execute initially. If contraversive gaze shifts are also more prevalent than ipsiversive gaze shifts, one would expect to find shorter saccade latencies for more eccentric orbital positions, on average (as we did in our data). Supporting this eye-head coordination account, individuals with a propensity for more head movement during head un-restrained gaze shifts also exhibit a more pronounced (larger slope) orbital position-latency relation (in head-restrained conditions), and vice versa [28, 29]. In summary, the needs of eye-head coordination may partially explain why fixation duration depends on orbital position in our data. However, this doesn’t explain why the eye ends up centered in the orbit following gaze shifts. And there appears to be an additional, synergistic reason for the orbital position-latency relation — namely, that the biomechanics and energetic efficiency of the oculomotor muscles facilitates/expedites re-centering of the eye in the orbit. This will be the focus on the next section.

### 3.6 Oculomotor biomechanics, energetic efficiency, and re-centering

The idea that the brain optimizes energetic costs has proven to be a useful predictive framework for understanding un-intuitive motor behaviors in walking [5]. Additionally, task- and motor-dependent cost functions in optimal feedback control models have proven successful in predicting behavior for other motor systems such as reaching [6, 35]. The same may be true for understanding oculomotor behavior like re-centering and the return phenomena we observed in the present study. Fixations at higher eccentricities require greater muscular force [36], which suggests that they should both be rarer and maintained for a shorter duration if the system is attempting to minimize energetic costs. Furthermore, the oculomotor muscle fibers, which are differentially recruited in the execution of large saccades, are much less fatigue-resistant than those that keep the eye in the primary position [37]. Re-centering of the eye in the orbit may also be facilitated by biomechanical and energetic imbalances between centripetal and centrifugal saccades (those that move the eye from an eccentric to a centered position in the orbit and the opposite, respectively). Centripetal saccades require more force to execute [38], simply due to the fact that passive elastic forces in the oculomotor muscles pull the eye to a centered position at all times, whereas larger, active forces are required to make a centrifugal saccade [39, 40]. Re-centering the eyes in the orbit may not only reduce motor costs at the present moment, but also in the near future — specifically it may reduce unnecessary head movements. A head movement is necessary to explore targets beyond the eye’s maximal position in the orbit, but head movements are more costly than eye movements, both in total energy expended and in opportunity costs incurred because of their slow speed of execution (versus ocular saccades) relative to the fast speed of internal processes like perception and planning. Additionally, orbit-eccentric eye orientations may increase the stochasticity of the visuomotor reference frame transformation required to generate actions from those processes, leading to inaccurate or delayed movement execution [41–44]. Facing an uncertain future, a centered eye position allows for exploration of a larger range of targets with less costly ocular saccades in either direction, versus more costly head movements [31]. Energetic efficiency at the level of neural control of the musculature or more central aspects of fixation/saccade control may also contribute to the re-centering bias [31, 40]. Ocular discomfort and fatigue, signals used by the body to prevent injury [45], may also play a role.

### 3.7 Conclusion

In this paper, we argue that the tendency to re-center the eyes in the orbit gives rise to a number of interesting properties of real-world scanpaths, including the predominance of return saccades [16]. The re-centering tendency itself seems to be due to a combination of multiple, synergistic factors: (1) neural and muscular energetic efficiency, (2) the limits and needs of eye-head coordination, and (3) higher-level perceptual, attentive, planning, and inferential processes that attempt to identify and orient to future targets. We can only speculate about the importance of each factor. Our current findings cannot speak to the relative contributions of each and how these vary with task demands and environment. The development of models which quantify the contributions of these factors is critical to understanding fixation selection. Much of the literature on inhibition-of-return and re-centering has treated peripheral muscular constraints as an uninteresting null hypothesis, so it’s unclear how much of seemingly high-level eye movement behaviors can be explained by low-level factors. Given that it is far more tractable to model the biomechanics and energetics of the oculomotor muscles than to model perception and cognition, we argue that the former should be done first. As a simplest next step, existing biophysical models of the eye plant [36, 38] can be re-configured as scanpath models by combining them with probabilistic sampling and optimizers that minimize physical energy (which some models already output). Some work already exists in trying to approximate user effort with oculomotor energy computed by eye plant models [46]. In general, models of gaze behavior at many levels of abstraction can incorporate oculomotor costs by considering fixation eccentricity as a proxy for effort (for head-fixed observers) and treating this as a (negative) reward, including active inference models of gaze behavior [3] and reinforcement learning models [4]. Deep learning models may have already learned this cost implicitly, hence why some machine learning researchers (correctly) try to factor out the predictiveness of the center bias on its own when evaluating saliency/gaze prediction deep learning models [47].

### 3.8 Limitations

Our data set, recording device, and analyses have a number of clear limitations. The Tobii Pro Glasses 2 is a mobile eye tracker and as such, when the observer moves more, data quality can be reduced. Likewise, fixation detection becomes more difficult when the observer is moving more, because of the predominance of counter-rolling of the eye during translational and rotational VOR, versus static fixations. Hence, tasks in which observers moved more (e.g., Walk and Navigation) had worse data quality, which may explain larger inter-observer variability in a number of our analyses for these tasks. One limitation of our recording device is that it does not allow us to precisely localize the center of the orbit for each eye (see Methods), which some other eye trackers can do. Another limitation is that we could not precisely estimate ocular saccades in world-centered coordinates, as many previous head-fixed lab studies have done. Separating out ocular saccades, counter-rolling of the eye during VOR, smooth pursuit, and static fixations during real-world conditions when an observer is moving is an open research problem and beyond the scope of this work [48]. Instead, because we primarily interested in the properties of scanpaths, static fixations, and their approximate orbital positions, we focused on identifying static fixations properly using well-accepted methodology (I-VT; see Methods for details) and we did this estimation in head-centered coordinates in order to assess approximate orbital position [16]. One issue with head-centered coordinates is that it introduces an ambiguity — a detected gaze shift can be due to either a head rotation (during VOR) or pure eye rotation, of opposite signs. As a sanity check, we repeated our analyses in gaze-in-body coordinates (i.e., counter-rotating gaze positions according to momentary head rotations derived from the onboard inertial measurement unit [IMU]), and we found similar results for all our main analyses — the same sort of spatial and temporal return phenomena and similar saccade metrics (see Supplementary Information). This demonstrates that possible artifactual influences of head movement on gaze-in-head fixation positions were minimal, which makes sense because head movements are much slower than ocular saccades and all of our analyses concerned brief, 3-fixation sequences. Furthermore, when we only analyzed time windows with minimal head movement (<3 ^*°*^/s angular speed), we observed similar center biases in fixation position and duration.

### 3.9 Open Questions

We are unable to explain why past studies observed longer fixations preceding return than forward saccades, whereas we observed the opposite. This apparent lack of temporal IOR could be a result of differences in the type of tasks used and/or the ability to move the head and body freely. For example, many past studies identifying temporal IOR used free-viewing or visual search tasks, which incentivize exploration of the visual image, a demand which may inhibit the natural tendency to re-center the eye in the orbit. So this account predicts that these past studies will show a smaller proportion of re-centering saccades (versus other types of saccades) than our study. Additionally, this past work implies that temporal IOR was present in our data but obscured by a stronger eccentricity-dependent effect of fixation duration. Another explanation is that the orbital position-latency relation is overall stronger in head-unrestrained versus head-restrained conditions because it has a greater utility in active eye-head coordination. That said, it is known that the orbital position-latency relation still holds even in head-restrained conditions [13] and furthermore, the neck muscles show activity related to eye-head coordination and saccade planning even when observers are head-restrained [49]. So this raises a worthwhile empirical question concerning the relative contributions of two putative components in different task contexts: an eccentricity-dependent one (the orbital position-latency relation) and an eccentricity-independent one (temporal IOR). Another interesting avenue for future work is quantitatively modelling the energy and opportunity costs involved in generating head and eye movements. If such a model is able to predict re-centering as an optimal strategy, that would be useful.

## 4 Materials and Methods

### 4.1 Participants

Thirty participants (ten internal employees and twenty external participants) took part in the study. All participants had normal or corrected-to-normal vision. Self-exclusion criteria for participants was anyone outside of an age range of 18-65 years old. Gender and age demographics were not recorded, so we cannot account for any age or gender-related differences between participants. Data were collected internally at Reality Labs, Meta Platforms Inc., in accord with the Declaration of Helsinki, and the study went through the organization’s research ethics review process (including independent Legal, Operations, Safety, and Program Manager approvals). All participants provided informed consent for their participation in the study.

### 4.2 Data collection

Each participant performed a variable subset of nine possible tasks, which ranged from approximately 5-20 minutes in duration. Participants wore the eye tracker during task performance, and in some cases operated the device themselves and in other cases were assisted by a research assistant. Tasks were selected to span a range of normal day-to-day activities and because these activities might benefit from augmentation. Some participants only performed one task, and others performed up to 6 different tasks, obviating systematic within-participants comparisons. The task descriptions and number of participants who performed each were as follows:

1. Driving (N=7). Participants drove to or from work.
2. Browsing (N = 30). Participants browsed the web with their mobile phone while seated.
3. Lego (N = 30). Participants built a lego set using instructions while seated at a table.
4. Walk (N=9). Participants walked outside on a forest path.
5. Desk (N=24). Participants worked on their computer at their desk.
6. Pokémon (N=18) Participants played an augmented reality mobile game (Pokémon GO) on their phone outside a building [50].
7. Restaurant (N=19). Participants ordered a meal on a tablet and spoke to a waiter as well. Note that this task was “acted” in a conference room with a confederate acting as a “waiter,” and an electronic tablet for a “menu”.
8. Grocery (N=20). Participants simulated searching for grocery items and getting them. Note that this was artificially done internally by having participants locate specified items in a micro-kitchen.
9. Navigation (N=10). Participants were tasked with finding a specific conference room in a research building and navigating there using a digital map on the wall and their phone.

Note that for each task (except driving), each participant conducted the task in a very similar environment.

### 4.3 Gaze measurement

Gaze was recorded using a Tobii Pro Glasses 2 mobile head-mounted eye tracker. The eye tracker contains two eye cameras and eight infrared illuminators per eye, and uses a combination of corneal reflection, dark pupil, and stereo geometry to achieve gaze measurements at 50 Hz with an angular error of ±0.62^*°*^ on average [51]. The Tobii firmware also accounts for slippage using a 3D geometric eye model. The trackable gaze range of the eye camera was 80^*°*^ horizontally by 50^*°*^ vertically. Gaze data were stored and processed on-device by Tobii’s proprietary firmware. We will refer to the coordinate system of the gaze data as “gaze-in-head” — as the gaze measurements are always relative to the orientation of the head. Specifically, the 3D origin of the gaze-in-head data was internally computed by the Tobii Pro firmware based on the estimated eye locations relative to the eye cameras at the time of calibration. This origin is located at the midpoint of the inter-ocular axis on the surface of the Tobii headset. Therefore, gaze measurements from the two eyes were transformed such that they were with respect to a hypothetical cyclopean observer.

Participants followed a single-point calibration method supplied by the manufacturer [52]. During this calibration, participants held a card with a target at arm’s length and approximately at the interocular axis elevation. The system used this calibration sequence to determine the best fit eye model and where it was relative to the cyclopean origin based on their proprietary internal algorithm.

### 4.4 Fixation detection

To detect gaze fixations, we used Tobii’s I-VT fixation detection method (implemented in the Tobii Pro Lab software), which employs a velocity threshold of 30^*°*^ /s. This method attempts to exclude smooth pursuit and ocular following eye movements, which are also types of fixations, both because of our concerns about the gaze estimate quality during these eye movement and because the objects’ visual location in the head-centered camera coordinates change during these eye movements, which should be detrimental to our predictability calculations. Instead, we detect only “stable fixations.” The I-VT method also merges fixations that are within a short interval of time, and accounts for measurement noise in the gaze estimates inherent to the Tobii Pro Glasses 2 mobile eye tracker.

Mobile eye trackers produce gaze and fixation measurements that are similar in accuracy from those from a standard EyeLink device (SR Research, Ottawa, Ontario), particularly when analyzed on average across measurements and participants [48]. Furthermore, saccade amplitudes and fixation durations we observed were within the range of what others have obtained with more precise devices. Therefore, we were confident in the fidelity of the gaze measurements.

### 4.5 Saccade estimation

We estimated saccades simply as the vector of displacement between two subsequent estimated fixation points in gaze-in-head coordinates. As a result, what we call a “saccade” in our analyses is actually a gaze shift (i.e., some combination of an ocular saccade and counter-rolling of the eye during head movement and VOR). And we described this saccade / gaze shift in head-centered coordinates, that is, there is an ambiguity of whether an estimated eye position change was caused by a head or eye movement. Note that this wasn’t a concern because we were primarily interested in the properties of fixations and scanpaths, not saccades, and in particular the orbital position of the eye during fixation. That said, to determine whether these ambiguities could be having a large influence on our primary results, we analyzed saccade directions and amplitudes using our data and found that were comparable to known metrics in the literature (see Supplementary Information). As a sanity check, we also transformed the gaze-in-head data into gaze-in-body coordinates by counter-rotating the estimated eye position by the momentary head rotation. The obtained saccade metrics were similar, demonstrating that the influence of head movement and counter-rolling of the eye on the estimated gaze shifts was minimal relative to the influence of ocular saccades (see Supplementary Information).

### 4.6 Model

We used a random sampling model to generate synthetic scanpaths, which consisted of a sequence of N fixations, each comprising (1) a horizontal position in degrees relative to the head orientation, (2) a vertical position in degrees relative to the head orientation, and (3) a duration in seconds. To generate a synthetic scanpath, N random independent samples were drawn from the empirical probability distribution of fixation locations, where N is equal to the number of datapoints in the empirical distribution, generating horizontal and vertical positions for each synthetic fixation. Next, the duration of a synthetic fixation was determined by simply looking up the duration in the corresponding horizontal and vertical position in the empirical map of fixation durations. The empirical probability distribution of fixation locations and the duration map shared the spatial sampling rate, allowing for this “look-up table” approach. We synthesized scanpaths in this manner for each task separately (based on the task-specific distributions) and for the pooled data across all tasks (based on the pooled distributions).

The model has two primary assumptions: (1) the duration of a fixation is only determined by its position relative to the head orientation (according to the functional form of the average fixation duration map), and (2) each fixation is independent of the specific content of the environment and the past history of fixations. Regarding point (1), the probability distribution of fixation duration at each location in the spatial map was summarized with the mean, so the model also ignores this variability. This approach also specifically ignores any other factors that are known to shape saccade selection and generation. Note that the model is simple by design. It is not meant to reproduce many aspects of fixation and saccade behavior, but rather to probe how far one can get in explaining temporal dependencies in fixation position and duration without explicitly specifying any cognitive or perceptual processes, and by assuming a random, temporally independent process. In this way, what is interesting is, quantitatively, how much the model can and cannot explain — which gives clues about the relative contributions of the components that give rise to the gaze behavior (which includes cognition and perception).

## 5. Supplementary Information

### 5.1 Saccade direction and amplitude depended strongly on task

Saccade directions were strongly task-dependent (Fig. S9A). Some tasks showed a strong horizontal plane bias as has been observed in some previous studies of saccade statistics during free-viewing [17, 53]. Others showed an almost even mix of each cardinal direction, such that vertical saccades were as common as horizontal. This was likely driven by reading behavior in the browsing and restaurant task, when observers read text on a tablet or phone and scrolled. In other cases (Grocery, Lego), the distribution was more rounded, with more oblique angle saccades than in the other tasks.

Average saccade amplitudes ranged from 5-10^*°*^ between tasks (Fig. S9B). This is consistent with past findings that saccade amplitudes tend to be under 10^*°*^ on average. Saccades were smallest in the browsing task and largest in the grocery task. In the browsing task, participants mostly read text on their phone from a distance, generating small saccades. In the grocery task, participants often searched drawers while standing, such that their heads were angled down, then looked up and searched the room for other items, generating large saccades coupled with head rotation. We performed two-sided permutation tests on every pair of tasks, and 34/36 comparisons were significant following false discovery rate (FDR) correction. That is, nearly all differences in amplitude observed between tasks were statistically significant.

### 5.2 Possible artifactual influence of head movement on gaze position does not explain away our findings

Gaze shifts measured in gaze-in-head coordinates have a fundamental ambiguity: a change in estimated gaze position can be due to an ocular saccade or to a head movement due to counter-rolling of the eye during VOR. This creates the possibility for a potential artifact A situation in which — (1) the eye starts in the primary position fixated an object X, (2) the observer makes an ocular saccade to an object Y, and (3) then orients their head to object Y (centering the eyes in the orbit again) — will be registered as a return saccade, even though it’s not. To control for this possibility, we repeated all our analyses after counter-rotating the gaze-in-head estimates by the momentary head rotation, i.e., transforming them into gaze-in-body coordinates (which is equivalent to world-locked coordinates when observer translation is zero). We found that gaze shift amplitudes were similar in gaze-in-head and gaze-in-body coordinates (Fig. S10A), demonstrating that gaze shifts were most often or mostly driven by ocular saccades, and influence of head movement was minimal. Critically, return saccades were almost equally probable before or after accounting for head movement (spatial IOR ratio = 2.44 vs. 2.22, before vs. after; Fig. S10B, Fig. S10C). This demonstrates that the possibly problematic scenario described earlier is not as common as the more obvious case of a first ocular saccade followed by a second, return ocular saccade, without a large net head orientation displacement in-between. This may also have to due with the fact that head movement during a gaze shift is much slower than ocular saccade execution, and the time window of the analysis is just three fixations long. We also observed similar, distinct three-fixation sequences of fixation duration for forward versus return saccades in gaze-in-body coordinates (Fig. S10D), and the same outcome for the distance-matched control analysis (Fig. S10E). The difference in fixation duration preceding a return versus forward saccade was smaller in gaze-in-body than gaze-in-head coordinates, but critically we still did not observe a pattern that resembles past studies (i.e., a pattern consistent with temporal IOR). To assess the impact of head movement on the statistics of fixation position and duration, we simply re-ran these analyses (in gaze-in-head coordinates), but only for fixations during which head movement was minimal (less than 3 ^*°*^/s angular speed, chosen as a threshold to both keep the amount of data high enough to limit sampling error and to minimize head movement). We observed center biases with the same overall shape (Fig. S10F, Fig. S10G), demonstrating that head movements do not explain away these effects either.

**Fig. S1.**
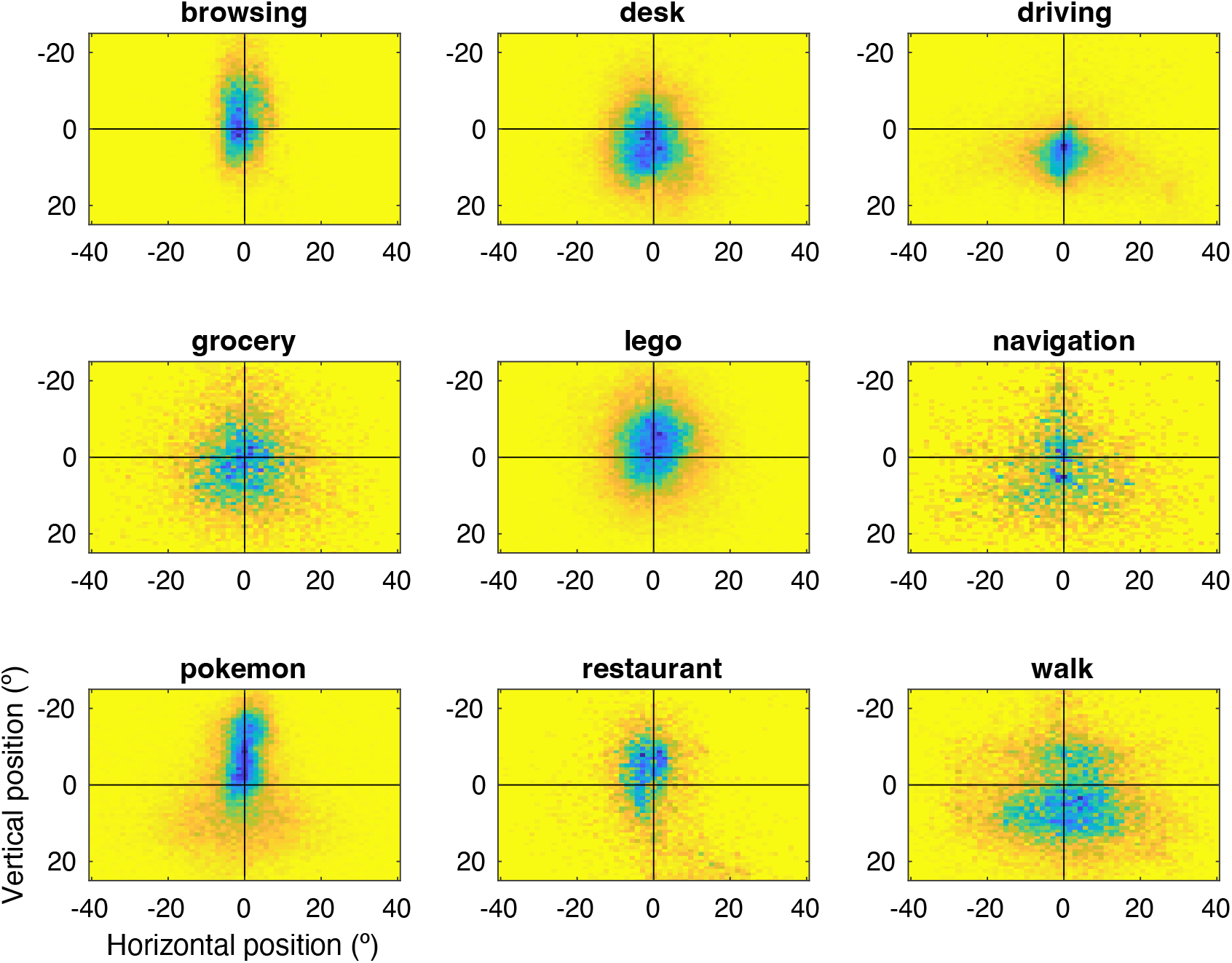
Probability distributions of fixation location for each task. Same format as Fig. 1, bottom panel. Colorbar limits were different for each subpanel, given that the amount of data varied between tasks.

**Fig. S2.**
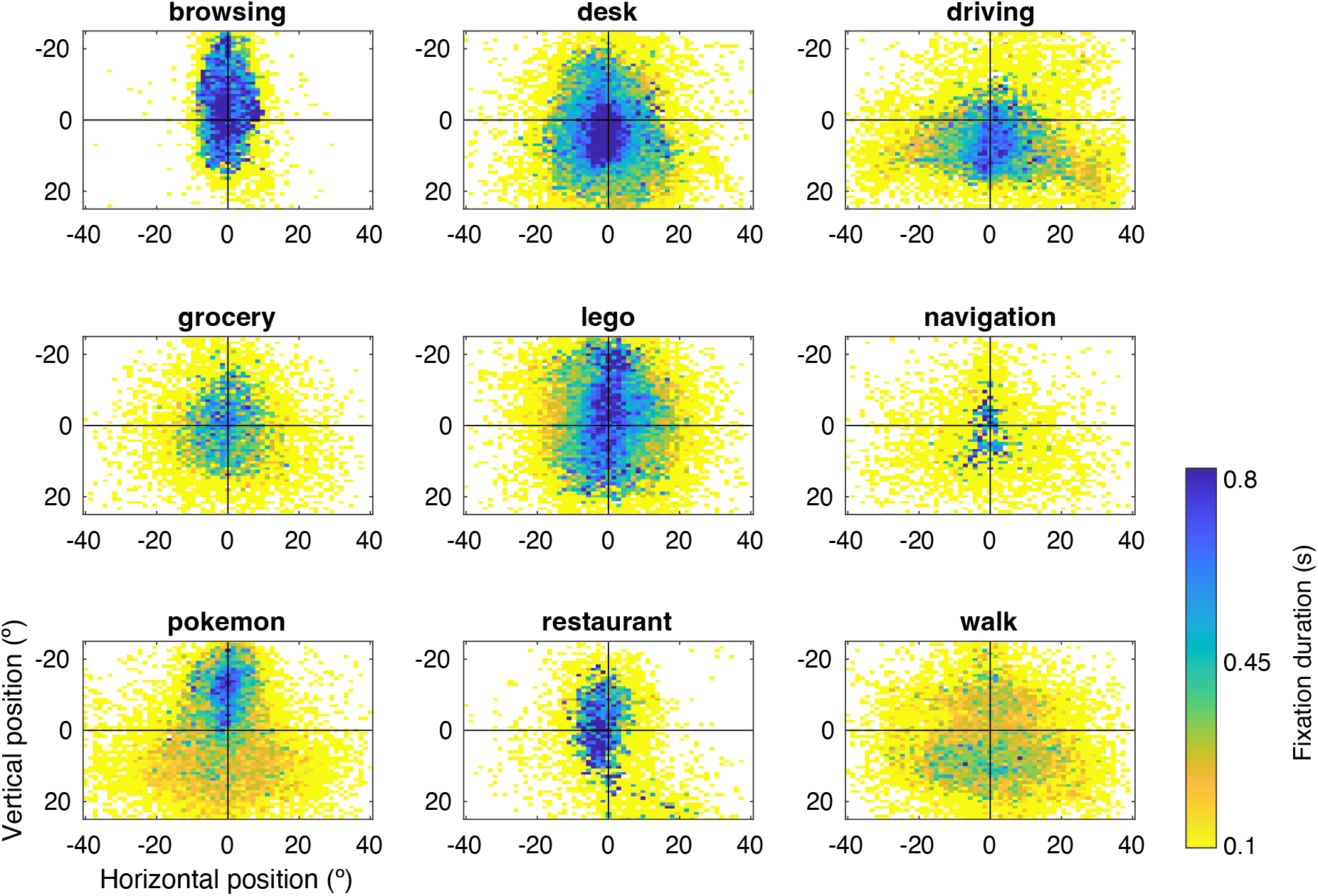
Spatial map of average fixation duration for each task. Same format as Fig. 1, middle panel. Colorbar limit is the same for each subpanel and was chosen to be consistent with Fig. 1. Note that some data points are above 0.8 s, for example in the browsing and desk tasks, but will be plotted at the darkest shade of blue available. White pixels are spatial bins for which there was only one data point / fixation or fewer.

**Fig. S3.**
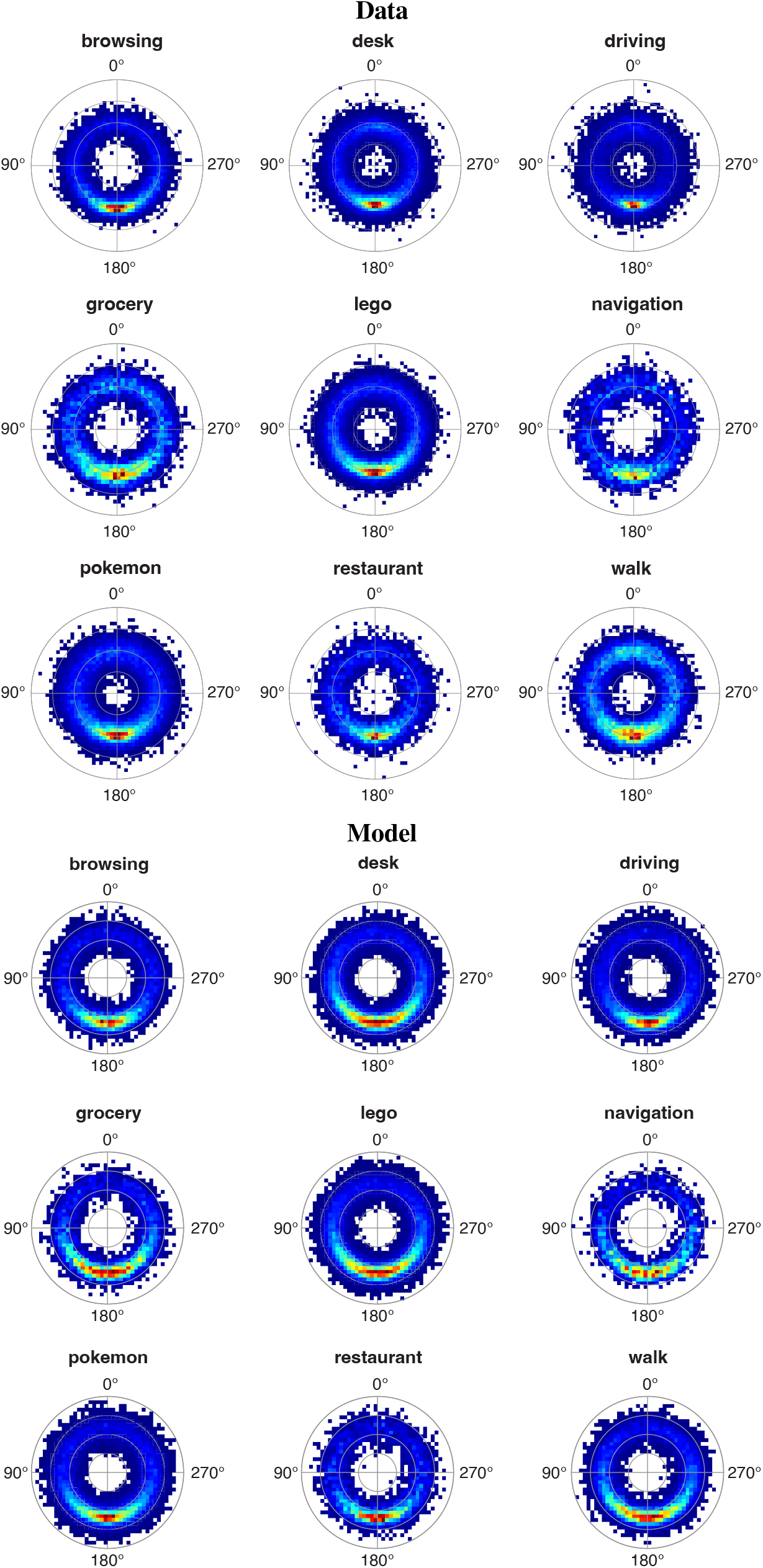
Relative saccade direction and amplitude joint distributions for each task, data versus model. Same format as Fig. 2 but for each task.

**Fig. S4.**
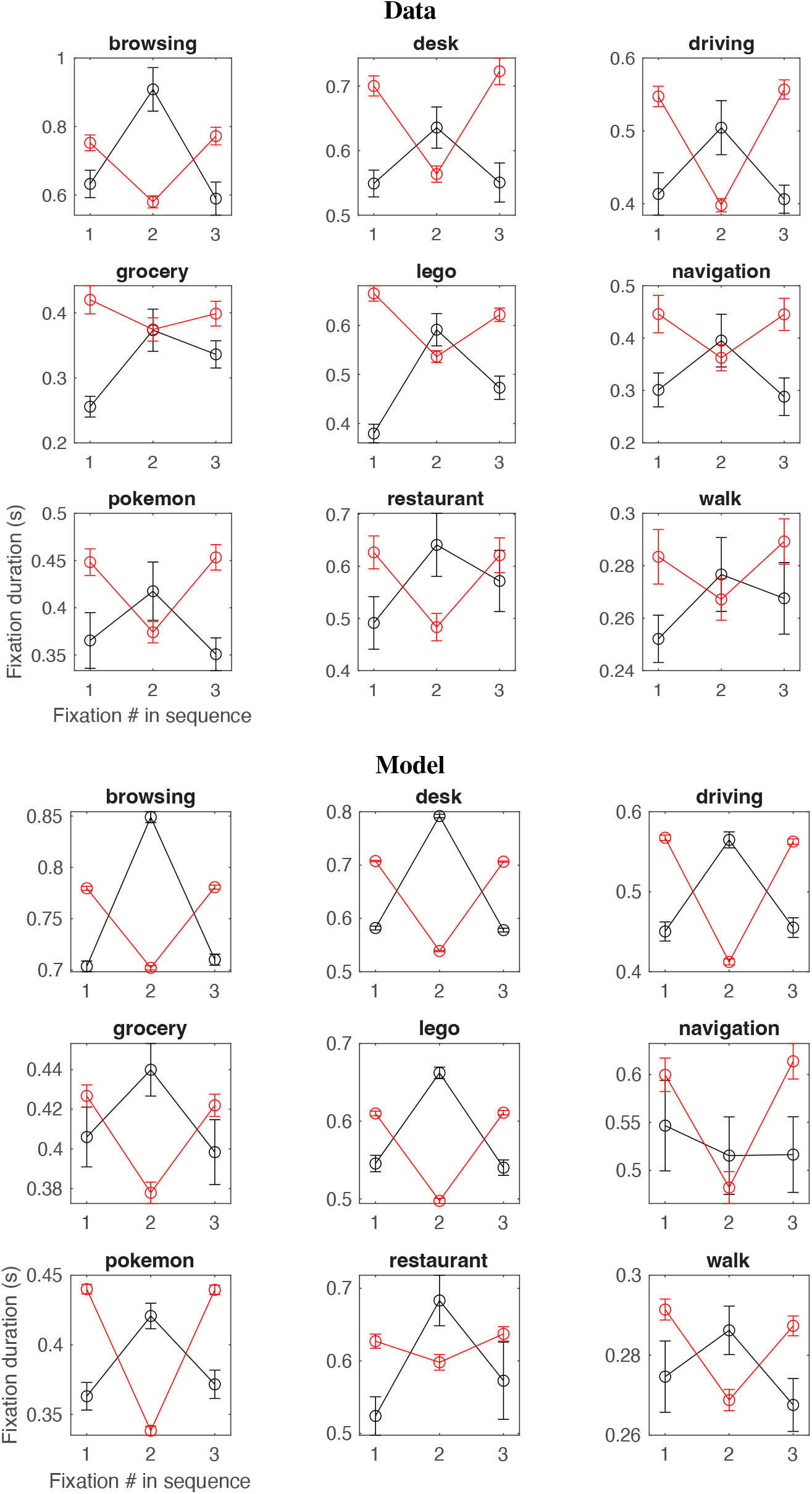
Fixation duration for three-fixation sequences for each task, data versus model. Same format as Fig. 3 but for each task.

**Fig. S5.**
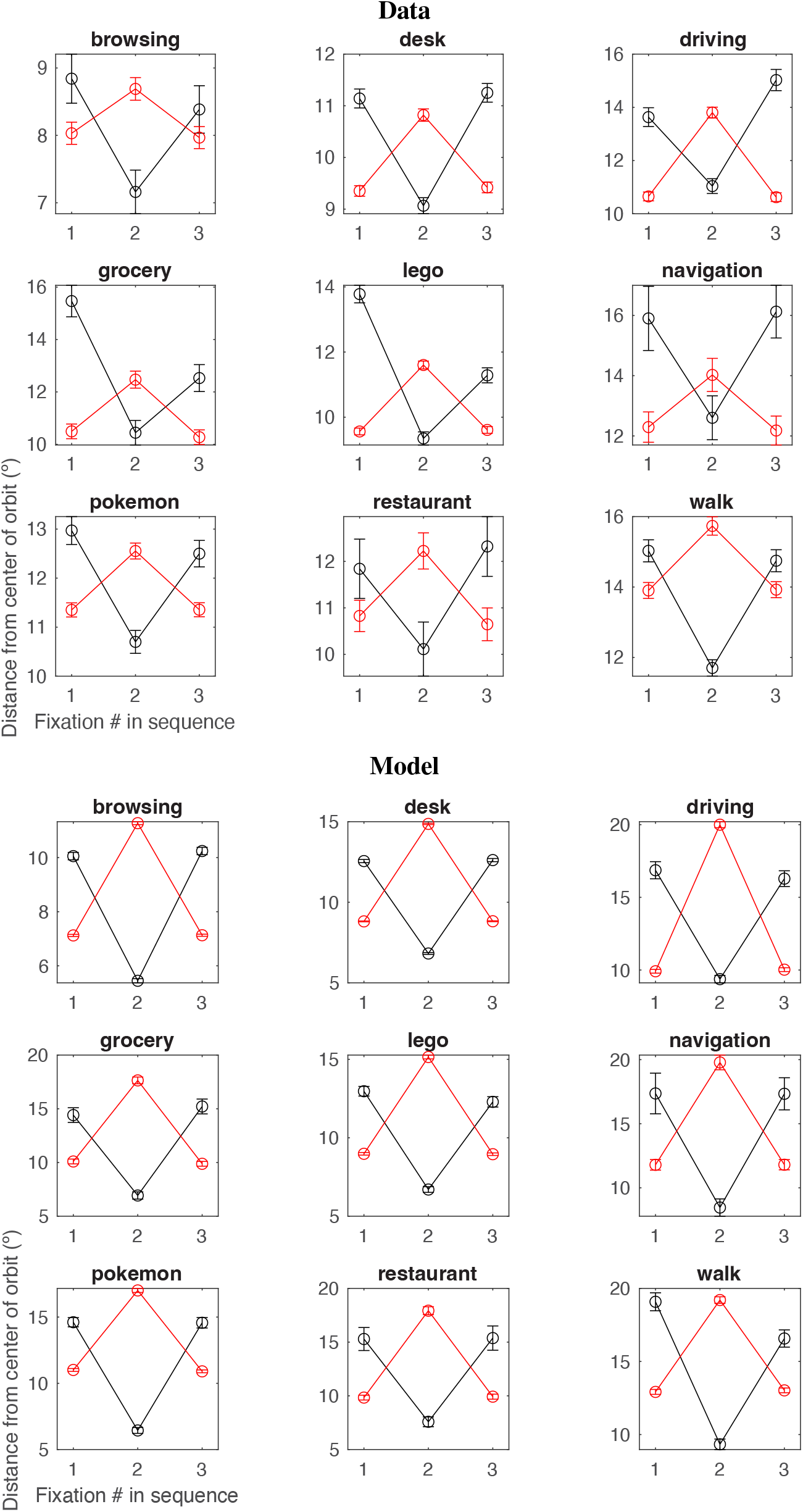
Distance from center of FOV for three-fixation sequences for each task, data versus model. Same format as Fig. 3 but for each task.

**Fig. S6.**
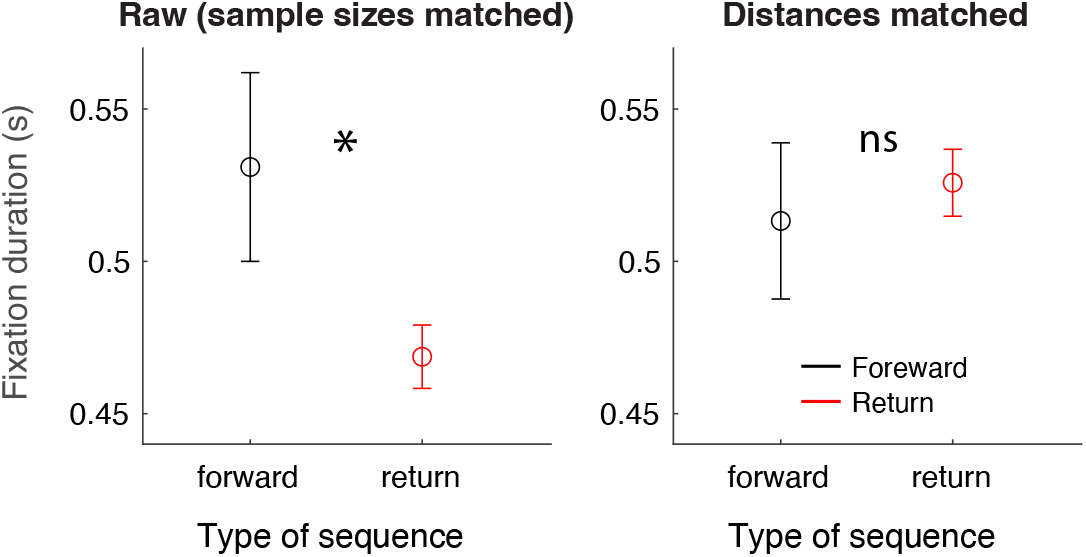
Control analysis: duration effects disappeared when distance from the center of FOV was fixed. Panel 1, average fixation duration in seconds preceding a forward or return saccade. Same data as shown in Fig. 3 but resampled such that sample sizes are equivalent in panels 1 and 2 (i.e., equivalent after distance thresholding has been applied). Shown is one example draw from the distributions that was significant (500 permutation tests, one for each resampling, median p-value = 0.0604, 46% of p-values were *<* 0.05, N permutations for each test = 1000). Panel 2, duration of fixation preceding forward or return saccade, after matching distance from the center of the FOV for forward and return sequences (permutation test, p = 0.67, N permutations = 1000). Matching consisted of finding sequences that were within 2^*°*^ of the median of the distribution of distances (i.e., where there was the most data). Distance matching was done for each participant and task separately. All plots depict pooled data across all recordings.

**Fig. S7.**
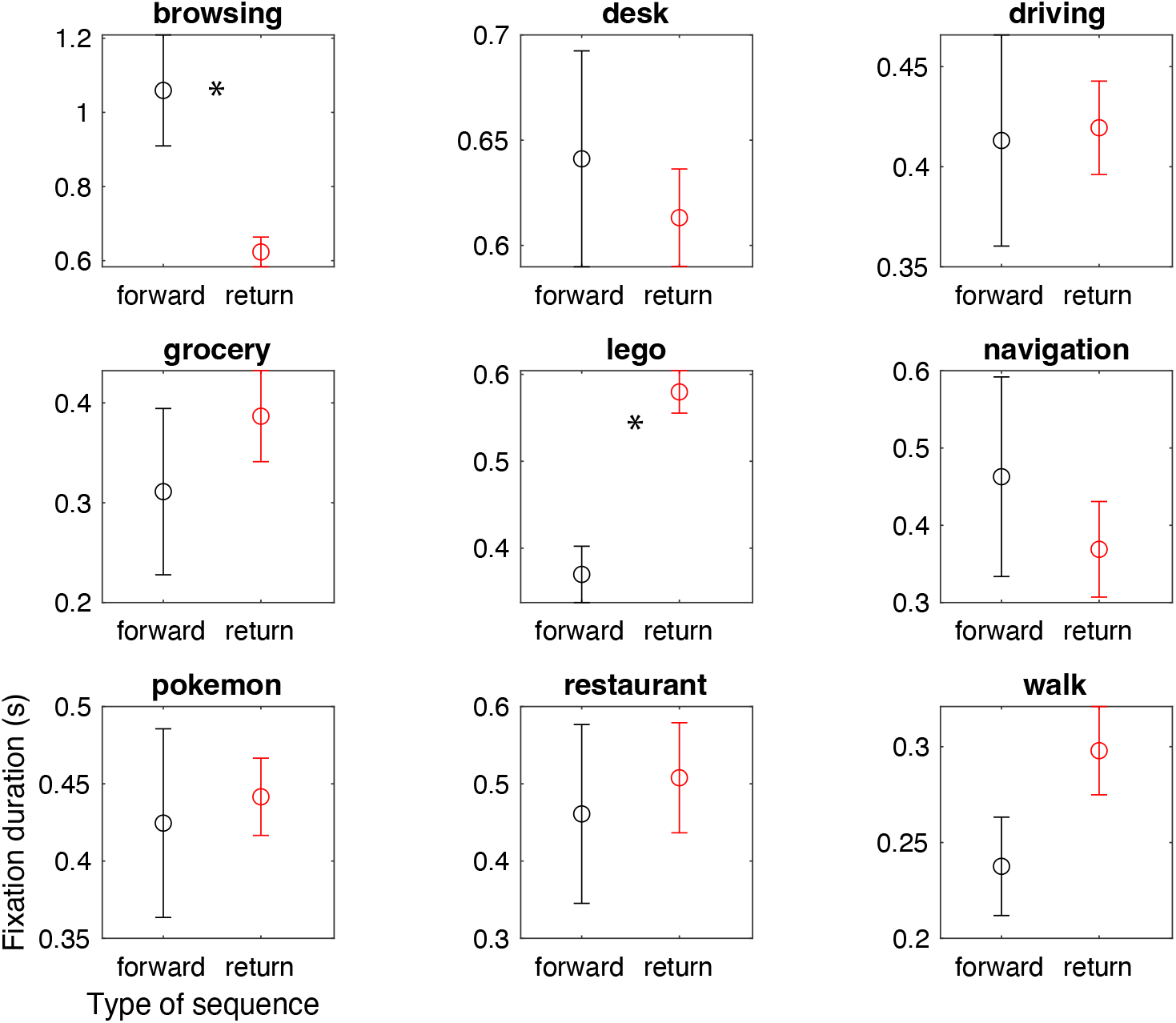
Control analysis for each task. Same format as Fig. S6, right subpanel, where distances were matched, except for each task separately. Asterisks represent outcome of two-tailed permutation tests, 1000 permutations each. P was *>* 0.05 for each task except browsing (p = 0.002) and lego (p = 0.001).

**Fig. S8.**
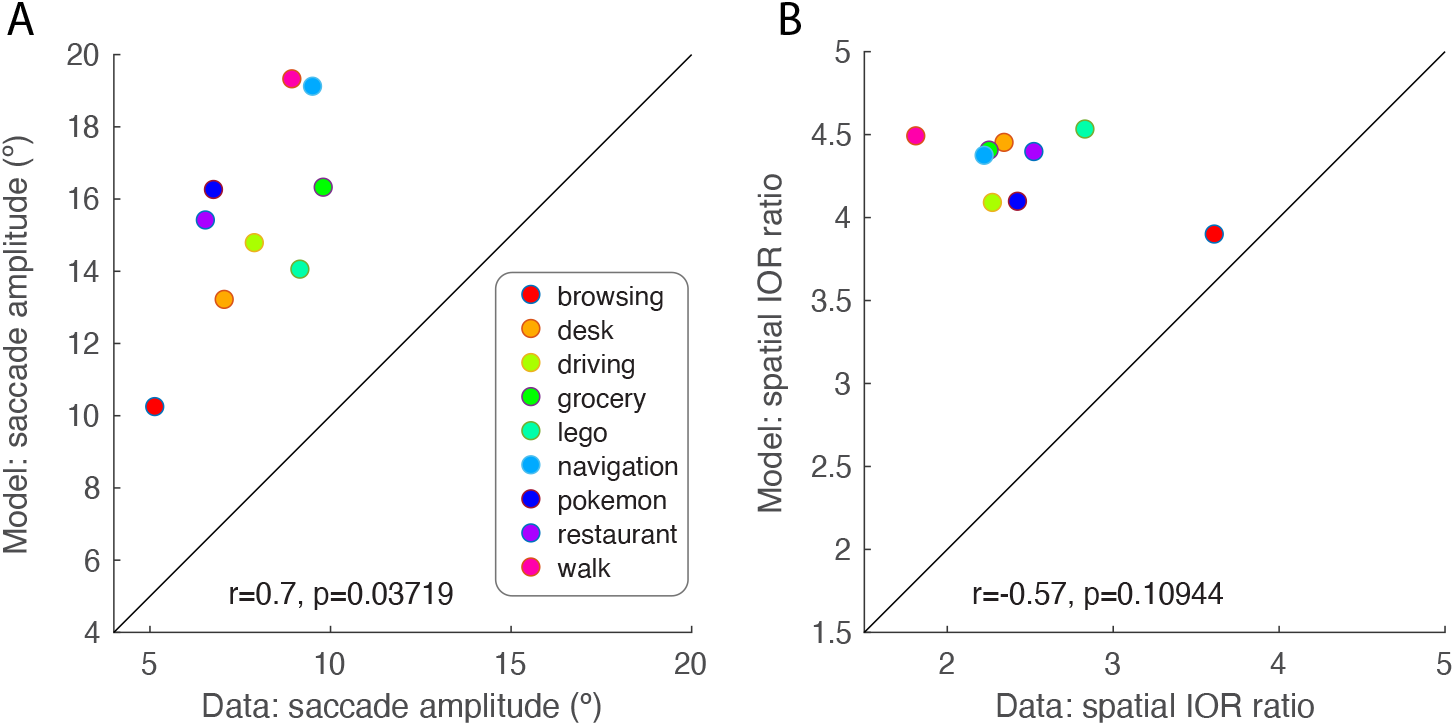
A random sampling model overestimates both saccade amplitude and the ratio of occurrence of return vs. forward saccades by around a factor of two. A. Median saccade amplitude (^*°*^) for the data versus the model, in each task. A model based on random sampling of the fixation density captures task dependency in saccade amplitude, hence the significant correlation, but overestimates amplitudes by around a factor of two. This overestimation is consistent with previous reports (Bays & Hussain, 2012), and demonstrates that people make smaller saccades than would be expected by the time-integrated statistics of their gaze locations. B. Spatial IOR ratio, the ratio of return to forward saccade occurrence, defined by slicing the relative saccade amplitude/direction joint distribution into four equal quadrants defined by the oblique angles and comparing the density in the two quadrants centered on precise return and forward saccades. The model overestimates this ratio by around a factor of two and doesn’t capture task dependency in the ratio.

**Fig. S9.**
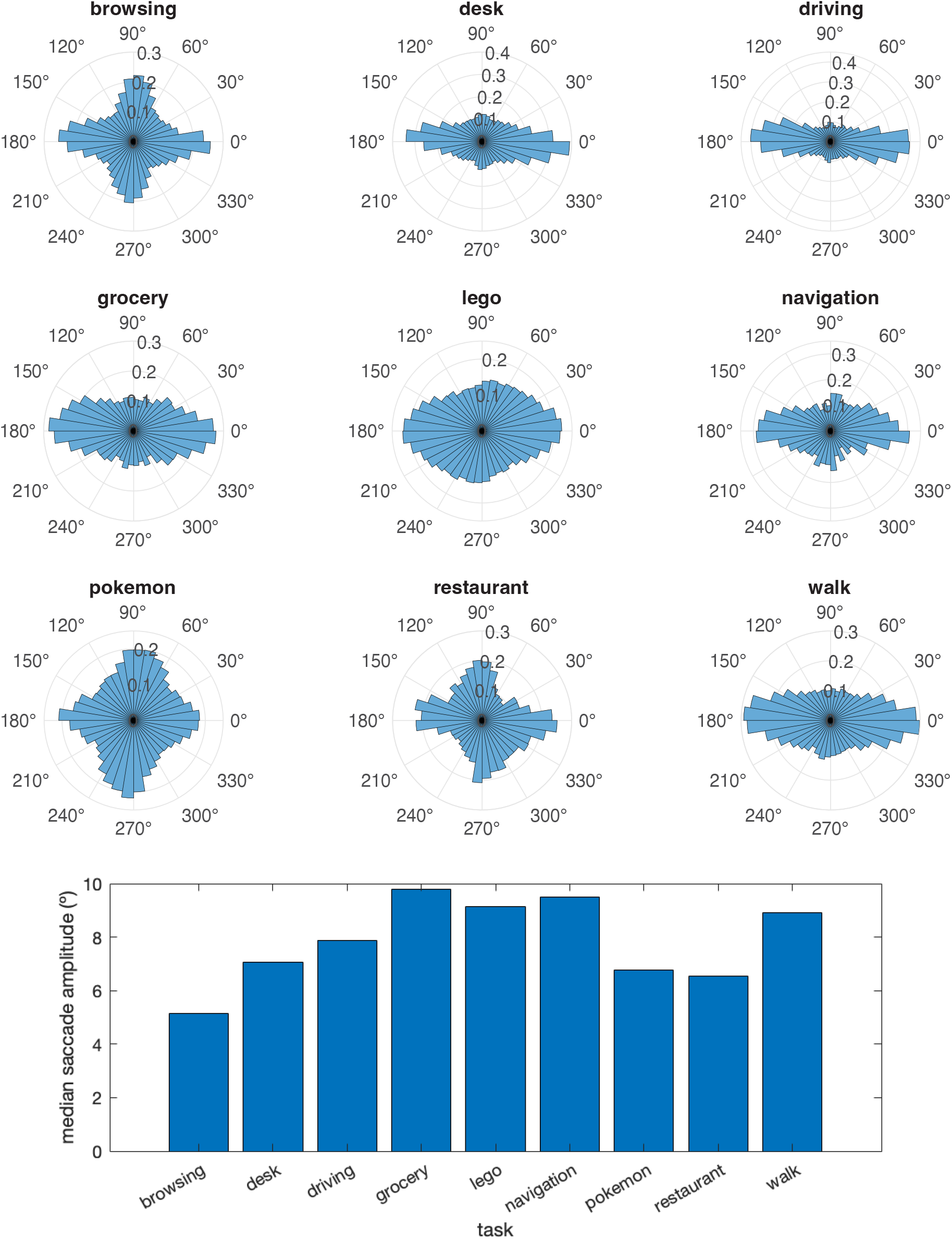
Saccade direction distributions and median amplitudes for each task. A. Saccade direction histograms for each task. Radius indicates probability and polar angle indicates saccade direction (where 90^*°*^ is vertical). B. Median saccade amplitude (^*°*^) for each task. Median is plotted instead of the full distributions because the distribution shape is similar for each task and quite broad so it’s difficult to tell where the center is by eye, even in log space.

**Fig. S10.**
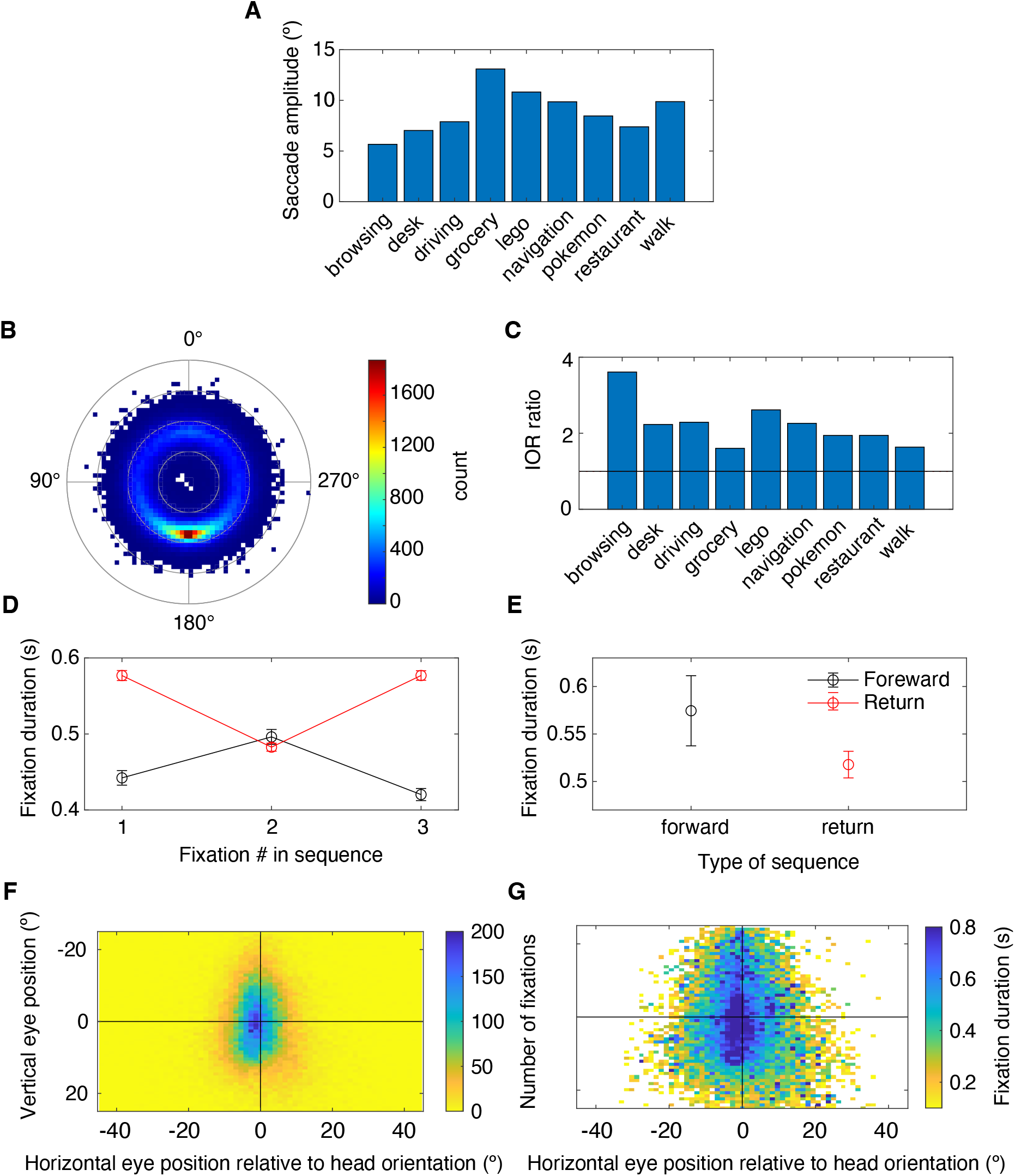
Possible artifactual influences of head movement on gaze position do not explain our findings. A-E, same format as prior figures but in gaze-in-body coordinates, i.e., after counter-rotating gaze-in-head positions by momentary head rotation. See prior figure captions for detailed explanations. A. Saccade amplitude (º). B. Relative saccade amplitude and directions. C. Spatial IOR ratio. D. Temporal sequences of fixation duration for forward and return saccades (same format as Fig 3). E. Distance-matched control analysis (same format as Fig. S6). F-G, same format as Fig 1, but only for fixations during which head movement was minimal (<3 ^*°*^/s angular speed). F. Probability distribution of fixation position. G. Spatial map of average fixation duration (note that only bins with greater than 2 data points are shown in color, to reduce sampling error while also showing rough shape of function).

## References

[1] Jiri Najemnik and Wilson S. Geisler. Eye movement statistics in humans are consistent with an optimal search strategy. Journal of vision, 8 3:4.1–14, 2008.

[2] Benjamin W Tatler, Mary M. Hayhoe, Michael Francis Land, and Dana H. Ballard. Eye guidance in natural vision: reinterpreting salience. Journal of vision, 11 5:5, 2011.

[3] M. Berk Mirza, Rick A Adams, Karl J. Friston, and Thomas Parr. Introducing a bayesian model of selective attention based on active inference. Scientific Reports, 9, 2019.

[4] Angela Radulescu, Bas van Opheusden, Frederick Callaway, Thomas L. Griffiths, and James M. Hillis. Modeling human eye movements during immersive visual search. bioRxiv, 2022.

[5] Geoffrey L. Brown, Nidhi Seethapathi, and Manoj Srinivasan. A unified energy-optimality criterion predicts human navigation paths and speeds. Proceedings of the National Academy of Sciences of the United States of America, 118, 2020.

[6] Stephen H. Scott. Optimal feedback control and the neural basis of volitional motor control. Nature Reviews Neuroscience, 5:532–546, 2004.

[7] Xuedong Wei and Alan A. Stocker. Lawful relation between perceptual bias and discriminability. Proceedings of the National Academy of Sciences, 114:10244 –10249, 2017.

[8] Judit Gervain and Maria Neimark Geffen. Efficient neural coding in auditory and speech perception. Trends in Neurosciences, 42:56–65, 2019.

[9] Rafael Polanía, Michael Woodford, and Christian C. Ruff. Efficient coding of subjective value. Nature neuro-science, 22:134 –142, 2018.

[10] Arthur Prat-Carrabin and Michael Woodford. Efficient coding of numbers explains decision bias and noise. Nature Human Behaviour, 6:1142–1152, 2022.

[11] Raymond M. Klein. Inhibition of return. Trends in Cognitive Sciences, 4(4):138–147, 2000.

[12] Laurent Itti and Christof Koch. Computational modelling of visual attention. Nature Reviews Neuroscience, 2:194–203, 2001.

[13] James H. Fuller. 101Comparison of Head Movement Strategies among Mammals. In The Head-Neck Sensory Motor System. Oxford University Press, 04 1992.

[14] Benjamin W. Tatler. The central fixation bias in scene viewing: Selecting an optimal viewing position independently of motor biases and image feature distributions. Journal of Vision, 7(14):4–4, 11 2007.

[15] Po-He Tseng, Ran Carmi, Ian G. M. Cameron, Douglas P. Munoz, and Laurent Itti. Quantifying center bias of observers in free viewing of dynamic natural scenes. Journal of vision, 9 7:4, 2009.

[16] Mengmi Zhang, Marcelo Armendáriz, Will Xiao, Olivia Rose, Katarina Nanna Filippa Bendtz, Margaret S. Livingstone, Carlos R. Ponce, and Gabriel Kreiman. Look twice: A generalist computational model predicts return fixations across tasks and species. PLOS Computational Biology, 18, 2021.

[17] Paul M. Bays and Masud Husain. Active inhibition and memory promote exploration and search of natural scenes. Journal of vision, 12 8, 2012.

[18] Tom Foulsham, Esther Walker, and Alan Kingstone. The where, what and when of gaze allocation in the lab and the natural environment. Vision Research, 51:1920–1931, 2011.

[19] Flora Ioannidou, Frouke Hermens, and Timothy L. Hodgson. The central bias in day-to-day viewing. Journal of Eye Movement Research, 9, 2016.

[20] Steven G. Luke, Joseph Schmidt, and John M. Henderson. Temporal oculomotor inhibition of return and spatial facilitation of return in a visual encoding task. Frontiers in Psychology, 4, 2013.

[21] Steven G. Luke, Tim J. Smith, Joseph Schmidt, and John M. Henderson. Dissociating temporal inhibition of return and saccadic momentum across multiple eye-movement tasks. Journal of vision, 14 14:9, 2014.

[22] Lars Oliver Martin Rothkegel, Hans Arne Trukenbrod, Heiko Herbert Schütt, Felix Wichmann, and Ralf Engbert. Influence of initial fixation position in scene viewing. Vision Research, 129:33–49, 2016.

[23] Jacob Hadnett-Hunter, George Nicolaou, Eamonn O’Neill, and Michael Proulx. The effect of task on visual attention in interactive virtual environments. ACM Trans. Appl. Percept., 16(3), 2019.

[24] Zhiming Hu, Andreas Bulling, Sheng Li, and Guoping Wang. Fixationnet: Forecasting eye fixations in task-oriented virtual environments. IEEE Transactions on Visualization and Computer Graphics, 27(5):2681–2690, 2021.

[25] Yin Li, Alireza Fathi, and James M. Rehg. Learning to predict gaze in egocentric video. 2013 IEEE International Conference on Computer Vision, pages 3216–3223, 2013.

[26] Kari S. Kretch and Karen E. Adolph. Active vision in passive locomotion: real-world free viewing in infants and adults. Developmental science, 18 5:736–50, 2015.

[27] Michael Francis Land. Predictable eye-head coordination during driving. Nature, 359:318–320, 1992.

[28] James H. Fuller. Head movement propensity. Experimental Brain Research, 92:152–164, 1992.

[29] James H. Fuller. Eye position and target amplitude effects on human visual saccadic latencies. Experimental Brain Research, 109:457–466, 1996.

[30] Douglas Blair Tweed. Visual-motor optimization in binocular control. Vision Research, 37:1939–1951, 1997.

[31] Martin Paré and Douglas P. Munoz. Expression of a re-centering bias in saccade regulation by superior colliculus neurons. Experimental Brain Research, 137:354–368, 2001.

[32] A. John van Opstal, Klaus Hepp, Y. Suzuki, and V. Henn. Influence of eye position on activity in monkey superior colliculus. Journal of neurophysiology, 74 4:1593–610, 1995.

[33] Daniela Zambarbieri, Giorgio Beltrami, and Maurizio Versino. Saccade latency toward auditory targets depends on the relative position of the sound source with respect to the eyes. Vision Research, 35:3305–3312, 1995.

[34] Ruth M. Krebs, Mircea Ariel Schoenfeld, Carsten Nicolas Boehler, Allen W. Song, and Marty G. Woldorff. The saccadic re-centering bias is associated with activity changes in the human superior colliculus. Frontiers in Human Neuroscience, 4, 2010.

[35] Reza Shadmehr, Maurice A. Smith, and John W. Krakauer. Error correction, sensory prediction, and adaptation in motor control. Annual review of neuroscience, 33:89–108, 2010.

[36] Oleg Komogortsev. Eye movement prediction by oculomotor plant modelling with Kalman filter. PhD thesis, Kent State University, 2007.

[37] R. John Leigh and David S. Zee. The neurology of eye movements. Contemporary neurology series; 90. Oxford University Press, Oxford, 5th edition. edition, 2015.

[38] Ansgar R. Koene and Casper J. Erkelens. Cause of kinematic differences during centrifugal and centripetal saccades. Vision Research, 42:1797–1808, 2002.

[39] David A. Robinson. Control of eye movements. Comprehensive Physiology, pages 1275–1320, 1981.

[40] Jérôme Tagu, Karine Doré-Mazars, Judith Vergne, Christelle Lemoine-Lardennois, and Dorine Vergilino-Perez. Recentering bias for temporal saccades only: Evidence from binocular recordings of eye movements. Journal of Vision, 18(1):10–10, 01 2018.

[41] J. K. Burns and Gunnar Blohm. Multi-sensory weights depend on contextual noise in reference frame transforma-tions. Frontiers in Human Neuroscience, 4, 2010.

[42] J. K. Burns, Joseph Y. Nashed, and Gunnar Blohm. Head roll influences perceived hand position. Journal of vision, 11 9, 2011.

[43] Hooman Alikhanian, Schubert Ribeiro de Carvalho, and Gunnar Blohm. Quantifying effects of stochasticity in reference frame transformations on posterior distributions. Frontiers in Computational Neuroscience, 9, 2015.

[44] T. Scott Murdison, Dominic I. Standage, Philippe Lefèvre, and Gunnar Blohm. Effector-dependent stochastic reference frame transformations alter decision-making. Journal of Vision, 22, 2022.

[45] Edward R. Perl. Ideas about pain, a historical view. Nature Reviews Neuroscience, 8:71–80, 2007.

[46] Dan E. Tamir, Oleg V. Komogortsev, and Carl J. Mueller. An effort and time based measure of usability. In WoSQ ‘08, 2008.

[47] Matthias Kümmerer, Thomas S. A. Wallis, and Matthias Bethge. Information-theoretic model comparison unifies saliency metrics. Proceedings of the National Academy of Sciences, 112:16054 –16059, 2015.

[48] Stefan Dowiasch, Peter Wolf, and Frank Bremmer. Quantitative comparison of a mobile and a stationary video-based eye-tracker. Behavior Research Methods, 52:667 –680, 2019.

[49] Debaleena Basu, Naveen Sendhilnathan, and Aditya Murthy. Neck muscle activity reflects neural patterns of sequential saccade planning in head-restrained primates. Journal of neurophysiology, 2022.

[50] Niantic Inc. and Nintendo / Creatures Inc. / GAME FREAK inc. Pokémon go. Mobile game, 2016.

[51] Tobii AB. Eye tracker data quality report: Accuracy, precision and detected gaze under optimal condi-tions—controlled environment. Tobii Pro Glasses 2 firmware v1.61., 2017.

[52] Tobii AB. Tobii Pro Glasses 2 User’s Manual. Tobii AB, 2016.

[53] Olivier Le Meur and Zhi Liu. Saccadic model of eye movements for free-viewing condition. Vision Research, 116:152–164, 2015.

